# A chromatogram-simplified *Streptomyces albus* host for heterologous production of natural products

**DOI:** 10.1101/612291

**Authors:** Asif Fazal, Divya Thankachan, Ellie Harris, Ryan F. Seipke

## Abstract

Cloning natural product biosynthetic gene clusters from cultured or uncultured sources and their subsequent expression by genetically tractable heterologous hosts is an essential strategy for the elucidation and characterisation of novel microbial natural products. The availability of suitable expression hosts is a critical aspect of this workflow. In this work, we mutagenised five endogenous biosynthetic gene clusters from *Streptomyces albus* S4, which reduced the complexity of chemical extracts generated from the strain and eliminated antifungal and antibacterial bioactivity. We showed that the resulting quintuple mutant can express foreign BGCs by heterologously producing actinorhodin, cinnamycin and prunustatin. We envisage that our strain will be a useful addition to the growing suite of heterologous expression hosts available for exploring microbial secondary metabolism.

## Introduction

The majority of clinically used antibiotics are derived from natural products produced by *Streptomyces* species and other closely related *Actinobacteria*, which were introduced into the clinic during a ‘golden era’ of antibiotic discovery spanning 1940-1960 [1]. The utility of these agents has been eroded over the last half-century due to misuse. As a consequence, there is now an urgent need to discover new antibiotics to treat bacterial infections, particularly those caused by drug-resistant ‘ESKAPE’ pathogens (*Enterococcus faecium, Staphylococcus aureus, Klebsiella pneumoniae, Acinetobacter baumanii, Pseudomonas aeruginosa*, and *Enterobacter species*) [2]. Growing concerns about resistance to antibacterial agents combined with the failure to find new leads from the screening of large libraries of synthetic compounds has led to a renewed interest in natural products discovery [3]. This renaissance has been fuelled to a large extent by the relatively inexpensive cost to sequence genomes of strains that produce promising bioactive small molecules. For example, there are currently >750 streptomycete genomes available in GenBank compared to four years ago when there were only ~150 [4]. Analysis of these genome sequences have illuminated the exciting prospect that the majority of microbial secondary metabolism has yet to be uncovered [5].

Further study of potentially interesting natural product biosynthetic pathways is often precluded by the poor growth characteristics of the strain and/or genetic intractability. Therefore, a heterologous expression strategy using a faster-growing and genetically tractable host is frequently adopted and indeed has become the method of choice for accessing new natural products and interrogating their biosynthesis. Technology for generating and screening large-insert genomic libraries (e.g. cosmid, BAC and PAC libraries) to obtain biosynthetic gene cluster (BGC) clones are well established [6,7]. These approaches are in the process of being superseded by recombination-based cloning methods for targeted ‘capture’ of DNA both from isogenic strains [8] and environmental DNA [9] as well as the ability to assemble natural/synthetic DNA *in vivo* and *in vitro* (recently reviewed by [10–12]).

Although fastidiously growing bacterial species, such as *Escherichia coli*, have been used for some heterologous expression studies, its use has thus far been rather limited because of poor expression of native promoter systems, but in some cases can be overcome by engineered promoter swaps either with the T7 promoter or those recognised by alternative RNA polymerase sigma factors [13,14]. As a consequence, streptomycetes and other related genera such as *Saccharopolyspora* and *Salinispora* are the go-to platform for heterologous expression, because they are metabolically robust with respect to availability of precursors and cofactors, and native promoter elements are more likely to be functional compared to alternative hosts [11,15]. Unsurprisingly, well studied species of *Streptomyces* such as *S. avermitilis, S. coelicolor, S. lividans* and *S. venezuelae* are commonly used for heterologous expression studies [16,17]. Several of the aforementioned species have been engineered for improved heterologous expression, principally via the mutation of endogenous BGCs [18–21].

Despite the utility of these strains, *S. albus* is still frequently used as a heterologous expression host. *S. albus* strains are seemingly distributed worldwide and encode a diverse range of natural products produced by a variety of biosynthetic systems [4,22]. They are genetically tractable and grow relatively quickly using conventional growth media and methodology. *S. albus* J1074 in particular has served as a workhorse for heterologous production of natural products over the last two decades and recently a strain with reduced biosynthetic pathway content was generated [23]. Relatively recently, we isolated a streptomycete from leafcutting ants that is closely related to *S. albus* J1074, which we designated *Streptomyces albus* S4 [24]. J1074 and S4 are phylogenetically closely related strains differing in only 12 nucleotide positions across >12 kb of DNA sequence representing 29 conserved single-copy conserved phylogenetic markers [22]. The two strains share ~80% of their BGCs, but S4 encodes a larger and more diverse complement of secondary metabolites [4]. The genetic tractability, robust and fastidious growth and ability to produce diverse secondary metabolites motivated us to modify this strain for easier detection of heterologously produced natural products.

There were two major aims of this work, the first aim was to construct a strain that when cultivated under routine conditions was unable to produce antifungal or antibacterial compounds, and the second aim was to construct a strain from which chemical extracts with reduced complexity could be generated. Here we describe the construction of a chromatogram-simplified strain of *S. albus* S4 where five BGCs have been mutated and we demonstrate its ability to heterologously produce actinorhodin, cinnamycin and prunustatin.

## Results and Discussion

### Construction of *Streptomyces albus* S4 Δ5

The *S. albus* S4 genome was previously sequenced [25,26] and it harbours at least 28 putative natural product biosynthetic gene clusters (BGCs) according to antiSMASH 3.0 (Fig. 1A, [27]. Chemical extracts prepared from *S. albus* S4 cultivated under a variety of culture conditions are dominated by antimycins and candicidin [24,26]. Therefore, abolishing production of these compounds was a top priority in our mutagenesis strategy. With the longer-term goal of ultimately developing an *S. albus* host as a platform to screen uncharacterised BGC clones for antibacterial activity, we also targeted the albaflavenone, surugamide and fredericamycin BGCs because these compounds have been reported to have antibacterial properties [28–30]. We mutated these five BGCs using conventional marker exchange mutagenesis (candicidin), PCR-targeted recombineering (antimycin) and CRISPR/Cas9 genome editing (albaflavenone, surugamide and fredericamycin). The workflow for this strategy is presented in Fig. 1B and is described in the materials and methods section. The final mutant strain, which we designated *S. albus* S4 Δ5, harbours a complete pathway deletion in the antimycin BGC, and deletions in key biosynthetic genes for the other four targeted BGCs. Deletions were first verified by PCR and the overall integrity of the *S. albus* S4 Δ5 strain was verified by genome sequencing using the Illumina MiSeq platform. The expected mutations were verified by mapping sequence reads to the *S. albus* S4 wild-type genome sequence [25]. A schematic representation of the relevant areas of the mapping are displayed in Fig. 1B and Figs. S1-S5.

**Fig 1.**
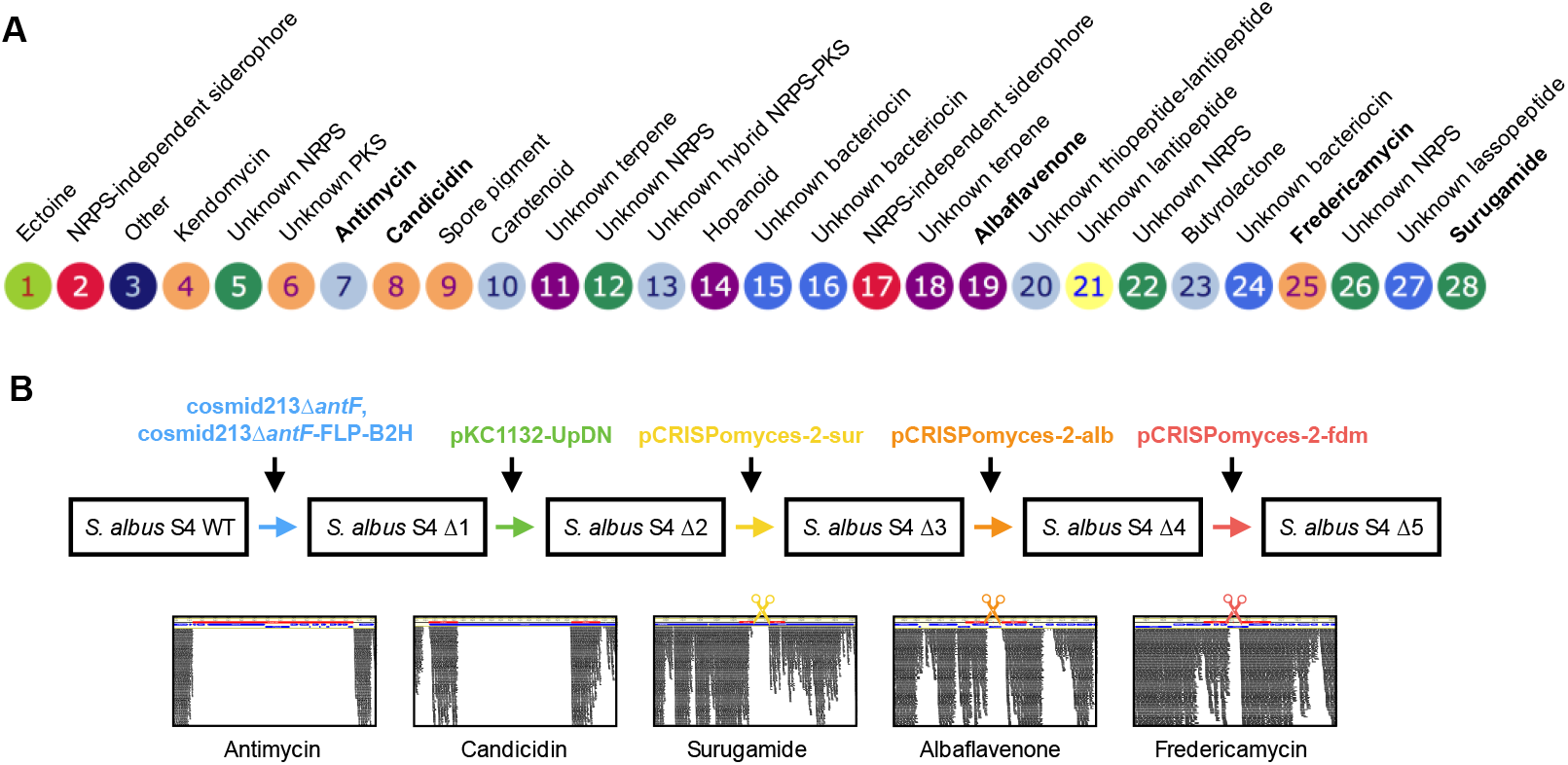
*Streptomyces albus* S4 biosynthetic gene clusters (BGCs) and their mutation. (A) Diagrammatic representation of putative BGCs identified by antiSMASH 3.0. The BGCs are numbered and the experimentally determined or bioinformatically deduced products are listed where known. BGCs targeted for mutagenesis in this study are indicated by bold text. (B) Schematic of the workflow used to create *S. albus* S4 Δ5. Knockout constructs are colour coded according to their targeted mutation and plasmid and strain designations are described in Table S1. The bottom panel depicts Illumina MiSeq reads mapped to the relevant locus of the *S. albus* S4 Δ5 genome illustrating that the desired mutation was achieved. The scissors indicate cleavage by the Cas9 nuclease. Full size versions of these images are shown in Figures S1-S5.

### Sporulation, biomass and bioactivity of Δ5

In order to ensure that serial passage (e.g. to cure plasmids with temperature-sensitive replicons such as that employed by pCRISPomyces-2) did not lead to mutation(s) adversely impacting sporulation and/or growth of Δ5, we assessed sporulation on MS agar and analysed its ability to produce biomass when cultivated using TSB. The Δ5 mutant strain sporulated equally as well as the wild-type strain, which indicated the absence of mutations deleterious to a normal developmental cycle and it also produced a comparable abundance of dispersed biomass to that of its parent (Fig. 2AB). Next, we analysed the bioactivity of ethyl acetate chemical extracts prepared from culture supernatants of *S. albus* S4 wild-type or Δ5 cultivated in liquid MS against *Candida albicans* and *Micrococcus luteus*, which revealed that extracts from the Δ5 strain were not bioactive (Fig. 2CD).

**Fig. 2.**
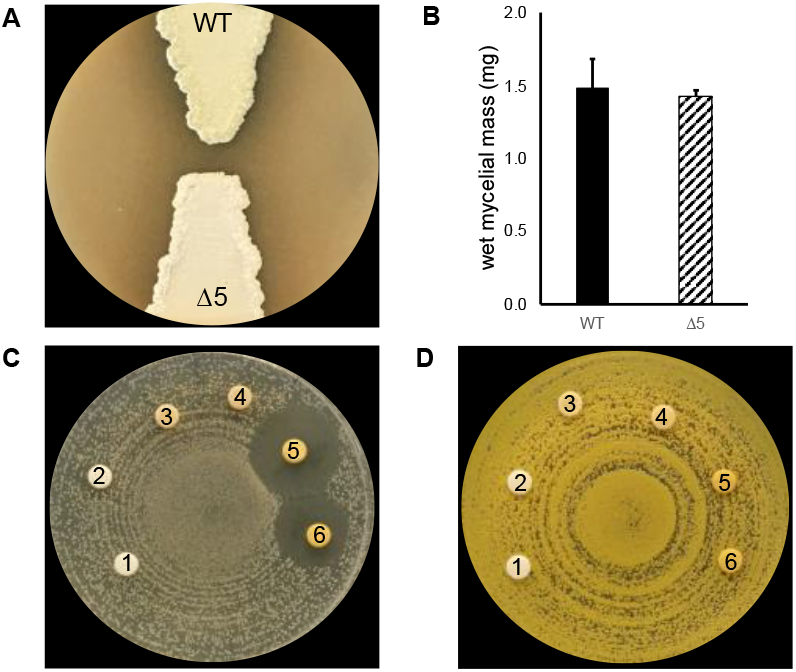
*S. albus* S4 Δ5 sporulates normally and produces biomass equal to that of the WT strain during growth in liquid media. (A) Sporulation of *S. albus* strains after growth on MS agar for 7 days. (B) Wet cell mass of *S. albus* strains originating from TSB cultures. The values reported are the means for a 20 ml sample of culture and the vertical bars represent the standard deviation (n = 3). The results are not statistically significantly different in a Student’s *t* test with a *P* value >0.60. Bioactivity of ethyl acetate chemical extracts generated from *S. albus* strains cultivated in liquid MS against (C) *Candida albicans* and (D) *Micrococcus luteus*. The discs on each plate are numbered and are as follows: 1, methanol; 2, extract from MS media; 3 and 4, *S. albus* Δ5 extract; 5 and 6, *S. albus* S4 WT extract. A zone of inhibited growth is apparent only for the wild-type extract against *C. albicans*.

### Chromatographic profile of Δ5

Chemical extracts generated from *S. albus* S4 wild-type are complex and chromatographs are dominated by candicidins and antimycins when using a standard aqueous/organic phase gradient of 5-95% acetonitrile or methanol [26,31]. In order to assess to what extent this complexity was reduced in extracts generated from the Δ5 mutant, we cultivated S4 wild-type and Δ5 using liquid MS and performed an ethyl acetate extraction of the resulting clarified culture supernatant. Next, extracts were analysed by HPLC, which as anticipated, revealed that the extracts generated from the Δ5 were far less complex than those generated from the wild-type strain (Fig. 3). The cleaner chromatographic background of the *S. albus* S4 Δ5 strain will simplify detection of heterologously produced natural products by HPLC or LC-MS and ease their subsequent purification.

**Fig. 3.**
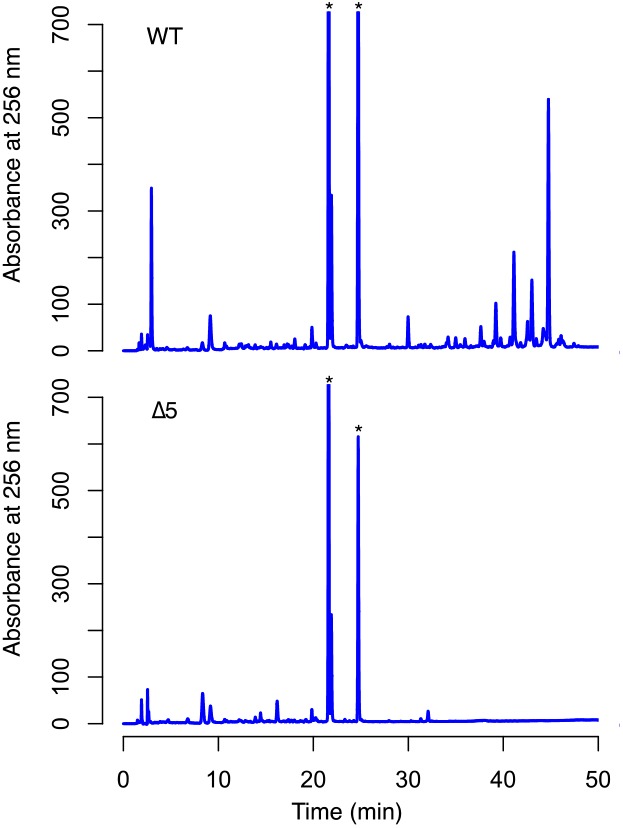
HPLC analysis of ethyl acetate extracts prepared from *S. albus* S4 WT and Δ5 strains. The HPLC chromatogram originating from the Δ5 strain is simpler in composition compared to that of the WT strain. The asterisks indicate HPLC peaks that are not fully visible in this image, but can be seen in Fig. S6.

### Heterologous production of actinorhodin, cinnamycin and prunustatin

As a proof of principle to demonstrate that S4 Δ5 can produce natural products encoded by other taxa, we introduced three foreign BGCs: the actinorhodin BGC from *S. coelicolor* A3(2), the cinnamycin BGC from *S. cinnamoneus* DSM 40646 and the neoantimycin/prunustatin BGC from *S. orinoci* B-NRRL 3379 [20,32,33]. These BGCs were selected in part due to accessibility, but also because they comprised three different biosynthetic systems. Actinorhodin is a type II polyketide antibacterial, cinnamycin is a ribosomally synthesised and post-translationally modified peptide antibacterial and neoantimycin/prunustatin is a hybrid non-ribosomal peptide / polyketide anti-cancer compound. Recombinant strains harbouring these BGCs were generated and cultivated using MS agar or TSB for seven days. Next, chemical extracts were prepared from the Δ5/Act, Δ5/Cin, Δ5/Prun strains and their parent and subsequently analysed by LC-HRMS (Fig. 4). Inspection of the resulting data revealed the presence of compounds consistent with ɣ-actinorhodin, cinnamycin and prunustatin in chemical extracts prepared from Δ5/Act, Δ5/Cin, Δ5/Prun, respectively and importantly, their absence from the Δ5 parental strain. These data indicate that *S. albus* S4 Δ5 is capable of heterologous expressing foreign BGCs from classes of varying biosynthetic logic.

**Fig. 4.**
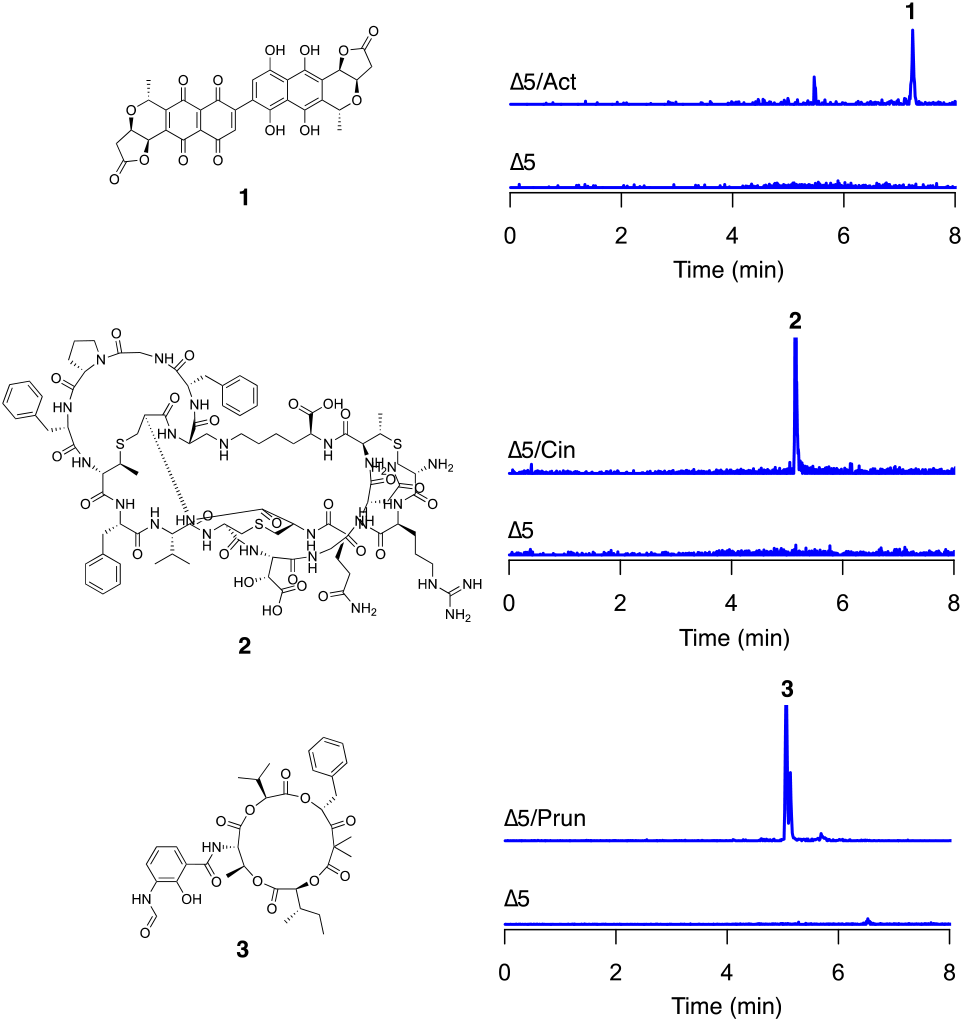
Heterologous production of ɣ-actinorhodin (**1**), cinnamycin (**2**) and prunustatin (**3**) by *S. albus* S4 Δ5. LC-HRMS analysis of chemical extracts prepared from the indicated strains. The *m/z* values corresponding to the [M+2H]^2+^ ions derived from ɣ-actinorhodin (C_32_H_22_O_14_), cinnamycin (C_89_H_125_N_25_O_25_S_3_), and prunustatin (C_36_H_44_N_2_O_12_) are shown. The intensity scale for extracted ion chromatograms is 1×10^4^ for ɣ-actinorhodin and cinnamycin, and 1×10^5^ for prunustatin.

### Summary and concluding perspectives

We have constructed a strain of *S. albus* S4 in which the antimycin, candicidin, albaflavenone, surugamide and fredericamycin BGCs have been mutated. We showed that the resulting strain, *S. albus* S4 Δ5, possessed a simpler chromatographic profile and did not possess antibacterial or antifungal activity using the growth media tested and also demonstrated that it could serve as a surrogate host for the production of actinorhodin, cinnamycin and prunustatin. Interestingly, during the course of our study we also tried to heterologously produce bicyclomycin and methylenomycin, but were unable to detect either metabolite by LC-HRMS. The unsuccessful attempt to heterologously produce bicyclomycin and methylenomycin underscores a key consideration for heterologous expression studies – host selection. For instance, *S. coelicolor* is a native producer of methylenomycin and was recently used as a host for bicyclomycin production [34], however it will not produce neoantimycin/prunustatin unless the biosynthetic genes are artificially expressed from constitutive promoters [33]. Indeed, there is no panacea in terms of a heterologous expression host, each one being somewhat enigmatic with peculiarities rooted in primary metabolism (i.e. precursor and co-factor supply) or promoter recognition elements. It is therefore advantageous and indeed wise when pursuing heterologous expression of a BGC to cast a wide net and screen multiple hosts. The *S. albus* S4 Δ5 strain constructed here will hopefully be a useful tool to that end.

## Materials and methods

### Growth media, strains and reagents

*Escherichia coli* strains were cultivated using Lennox agar (LA) or broth (LB) and *Streptomyces* strains were cultured using mannitol-soya flour (MS) agar, tryptic soy broth (TSB) or agar (TSA) [35]. Culture media was supplemented with antibiotics as required at the following concentrations: apramycin (50 μg/ml), carbenicillin (100 μg/ml), hygromycin (75 μg/ml), kanamycin (50 μg/ml), nalidixic acid (25 μg/ml). Enzymes were purchased from New England Biolabs and oligonucleotides were purchased from Integrated DNA Technologies. The bacterial strains, cosmids and plasmids used in this study are described in Table S1 and the oligonucleotides are described in Table S2.

### Mutagenesis of biosynthetic gene clusters

In order to delete the *ant* BGC, the *antF* gene on S4 Cosmid213 [31] was replaced with the apramycin resistance cassette from pIJ773 using PCR targeted mutagenesis as previously described [36]. The mutated cosmid was used to generate an apramycin resistant *S. albus* S4 *ΔantF* strain. Next, *antABCDEFGHIJKLMNO* on Cosmid213 were replaced with the apramycin resistance cassette in the same manner as above and subsequently removed by the Flp recombinase, after which the *bla* gene was replaced with the hygromycin resistance cassette also harbouring an *oriT* from pIJ10701 as previously described [36] to result in Cosmid213 ΔantFLP B2H. This cosmid was mobilised to *S. albus* S4 *ΔantF* and a single hygromycin resistant transconjugant was selected and subsequently passaged twice in the absence of selection prior to replica plate identification of an apramycin sensitive and hygromycin sensitive isolate that we named *S. albus* S4 Δ1. The integrity of this strain was confirmed by PCR using primers RFS236 and RFS237.

The *can* BGC was mutated using the previously constructed *fscC (STRS4 02234*) deletion plasmid, pKC1132-UpDn [26]. pKC1132-UpDn was mobilised to *S. albus* S4 Δ1 strain and a single apramycin resistant transconjugant was selected and passaged as above until an apramycin-sensitive isolate was identified, which we named *S. albus* S4 Δ2.

The surugamide, fredericamycin and albaflavenone BGCs were mutated using the pCRISPomyces-2 system describe previously [37]. First, a single-guide RNA protospacer was generated by annealing oligonucleotides EH_S9 and EH_S10 (surugamide), EH_S3 and EH_S4 (albaflavenone) and EH_S7 and EH_S8 (fredericamycin); the resulting DNA fragments were cloned into the BbsI site of pCRISPomyces-2 by Golden Gate Assembly. Second, two overlapping DNA fragments representing a homology-directed repair template were generated by PCR using the primers listed in Table S2. The overlapping PCR products were subsequently cloned into the XbaI site of protospacer-containing pCRISPomyces-2 plasmid using the NEBbuilder HiFi DNA assembly kit. The resulting CRISPR/Cas9 editing plasmids, pCRISPomyces-2-sur, pCRISPomyces-2-alb, pCRISPomyces-2-fdm were sequentially mobilised to *Streptomyces* and cured as described previously [38] and schematically illustrated in Figure S1 to result in the quintuple mutant strain described in this study, *S. albus* S4 Δ5.

### Genome sequencing and bioinformatics analysis

*S. albus* S4 Δ5 chromosomal DNA was sequenced by Microbes NG (Birmingham, UK) using the Illumina MiSeq platform. This resulted in the generation of 2,859,380 paired-end reads that were 250 nt in length. 2,809,652 of these reads were mapped the *S. albus* S4 wild-type chromosome (GenBank accession CADY00000000.1 [25]) using the Geneious assembler (version R8.1.19). Schematic representations of relevant loci depicted in Figs. S2-S6 were generated using Geneious version R8.1.19.

### Bioactivity assays

Indicator organism was cultivated overnight in LB at 37 °C. Overnight culture was diluted to an OD_625nm_ of 0.08 in LB and spread onto an LB agar plate using a rotary platform. Sterile paper discs (6 mm diameter) holding 60 μl of chemical extract were placed atop seeded indicator plates, which were subsequently incubated at 37 °C (*Micrococcus luteus*) or room temperature (*Candida albicans*) and visualized the following day.

### HPLC analysis

*S. albus* strains were cultivated in liquid MS (50 ml) whilst shaking at 220 rpm in a 250 ml flask at 30 °C for 7 days. Bacterial cells and solid matter from the culture medium were removed by centrifugation and 45 ml of supernatant were extracted with 90 ml of ethyl acetate. The extract was evaporated to dryness under reduced pressure and resuspended in 100% methanol (500 μl). The methanolic extract was centrifuged for 10 minutes at 16,000 x *g* to remove insoluble material prior to sample injection (10 μl) into a Dionex HPLC instrument. Compounds were separated on a Phenomenex Luna C18 column (5 μm, 150 x 4.6 mm) using the following gradient (solvent A: 5% acetonitrile, 0.1% formic acid, solvent B: 95% acetonitrile, 0.05% formic acid, flow rate 1 ml/min): 0-40 min, 0-100% B; 40-43 min, 100% B; 43-51 min 0% B.

### LCMS analysis

Clones of the actinorhodin, cinnamycin and neoantimycin BGCs were introduced into *S. albus* Δ5 by intergeneric conjugation from *Escherichia coli* ET12567/pUZ8002. Apramycin resistant transconjugants were verified to have received BGCs by PCR. For the production of actinorhodin and prunustatin, MS agar plates were seeded with either Δ5 or Δ5/Act or Δ5/Prun and cultivated for 7 days at 30 °C. The agar was then cut into small rectangular pieces and placed into an Erlenmeyer flask and metabolites extracted with ethyl acetate (100 ml) for 24 hrs. The ethyl acetate extract was decanted and concentrated *in vacuo* and the dried residue was resuspended in 100% methanol (500 μl). For the production of cinnamycin, *S. albus* S4 Δ5 and Δ5/Cin strains were cultured in 50 ml TSB for 7 days at 30 °C. The resulting mycelia was collected by centrifugation and extracted with 100% methanol (45 ml) for 4 hours. The methanolic extracts were evaporated to dryness *in vacuo* and resuspended in 100% methanol (500 μl) as above. Equal volumes of methanolic extract for three independently cultivated replicates of Δ5, Δ5/Act, Δ5/Cin, Δ5/Prun were pooled and centrifuged for 10 minutes at 16,000 x *g* to remove insoluble material prior to analysis by high resolution electrospray ionization liquid chromatography mass spectrometry (LC-HRMS). Only the supernatant (2 μl) was injected into a Bruker MaXis Impact TOF mass spectrometer and equipped with a Dionex Ultimate 3000 HPLC exactly as previously described [13].

## Acknowledgements

We are grateful to Iain Manfield for assistance with HPLC experiments, which were conducted in the University of Leeds Biomolecular Interactions Facility. We thank David Widdick and Mervyn Bibb for providing the cinnamycin and actinorhodin BGC clones, Andy Truman for providing the bicyclomycin BGC clone and Christophe Corre for providing the methylenomycin BGC clone. This work was supported by Biotechnology and Biological Sciences Research Council responsive mode grant BB/N007980/1 awarded to RFS. AF and DT were funded by PhD studentships funded by the University of Leeds.

## Ethical statement

No human or animal experiments were conducted during his study.

