## Supplementary Information for "A chromatogram-simplified *Streptomyces albus* host for heterologous production of natural products"

### Table of Contents

Supporting figures and tables

**Figure S1.** Illumina read mapping to the antimycin BGC in *S. albus* S4 Δ5.
**Figure S2.** Illumina read mapping to the candicidin BGC in *S. albus* S4 Δ5.
**Figure S3.** Illumina read mapping to the albaflavenone BGC in *S. albus* S4 Δ5.
**Figure S4.** Illumina read mapping to the surugamide BGC in *S. albus* S4 Δ5.
**Figure S5.** Illumina read mapping to the fredericamycin BGC in *S. albus* S4 Δ5.
**Figure S6.** Full scale HPLC trace for *S. albus* S4 WT and *S. albus* S4 Δ5.
**Table S1.** Bacterial strains, cosmids and plasmids.
**Table S2.** Oligonucleotide primers used in this study.

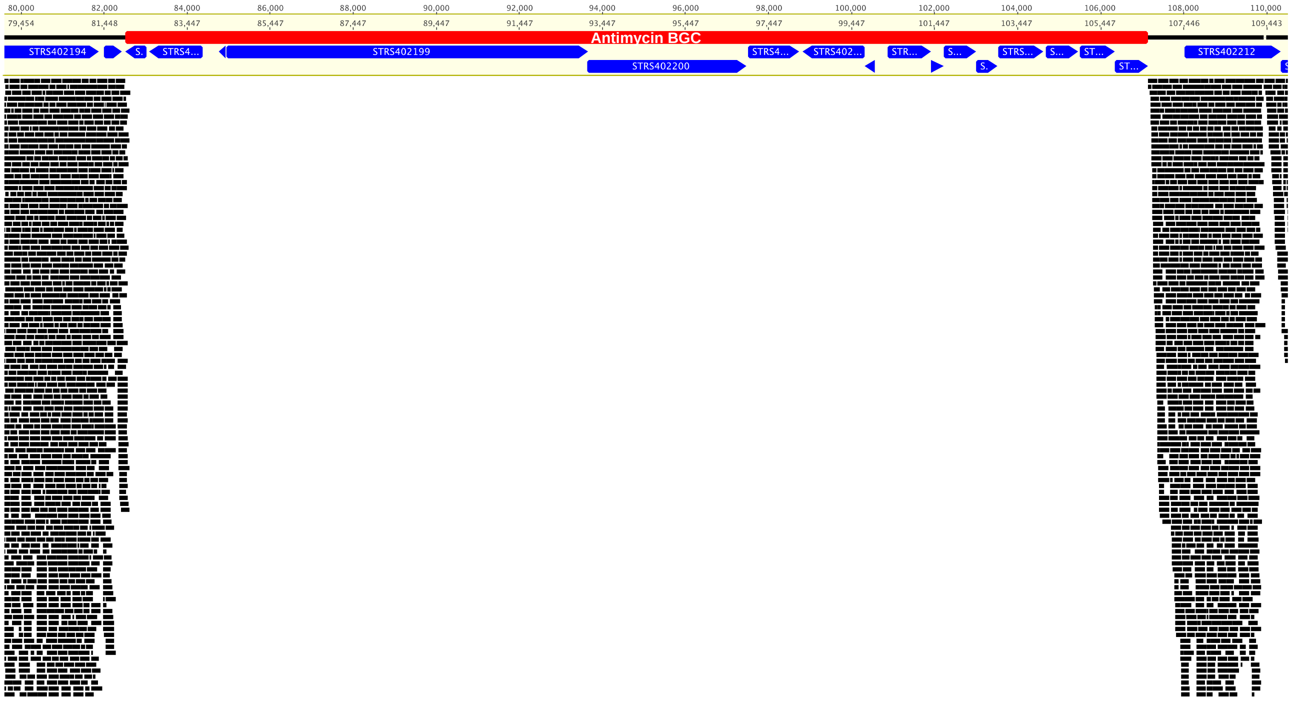


**Figure S1.** Deletion of the entire antimycin BGC in *Streptomyces albus* S4 Δ5. Black rectangles represent Illumina MiSeq reads mapped to the *S. albus* S4 Δ5 genome. The red rectangle indicates the antimycin BGC.

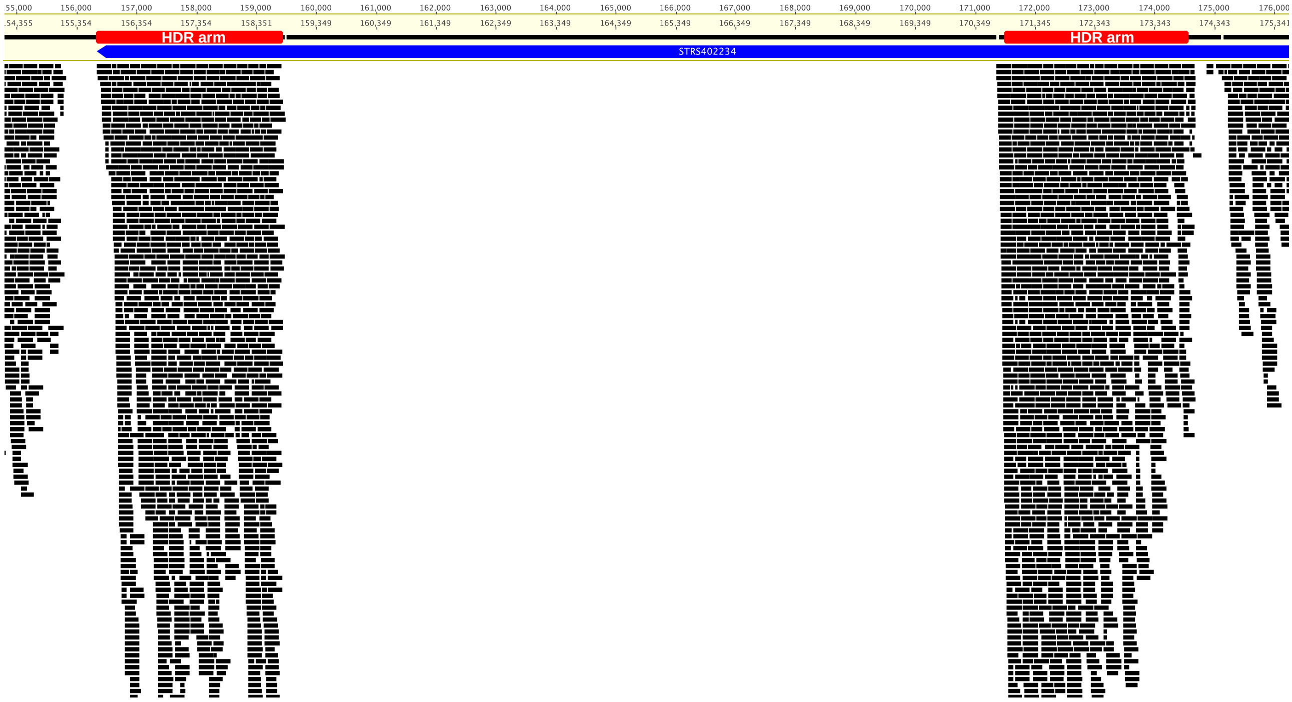


**Figure S2.** Mutagenesis of the candicidin BGC in *Streptomyces albus* S4 Δ5. Black rectangles represent Illumina MiSeq reads mapped to the *S. albus* S4 Δ5 genome. Red rectangles indicate the homology-directed repair arms used to delete the *STRS402234* gene (also known as *fscC*). HDR, homology-directed repair arm.

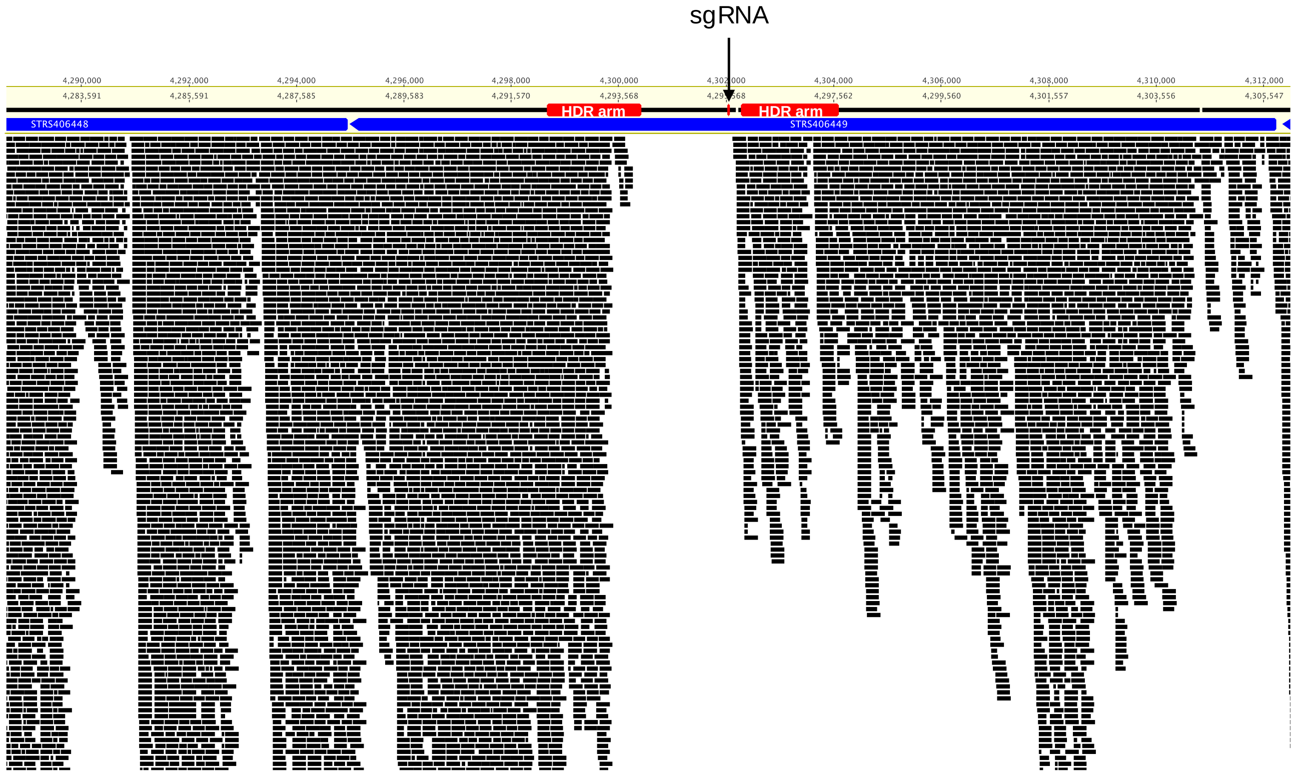


**Figure S3.** Mutagenesis of the surugamide BGC in *Streptomyces albus* S4 Δ5. Black rectangles represent Illumina MiSeq reads mapped to the *S. albus* S4 Δ5 genome. Red rectangles indicate the homology-directed repair arms used to repair the double strand break generated by the Cas9 targeting the *STRS406449* gene. HDR, homology-directed repair arm; sgRNA, single guide RNA.

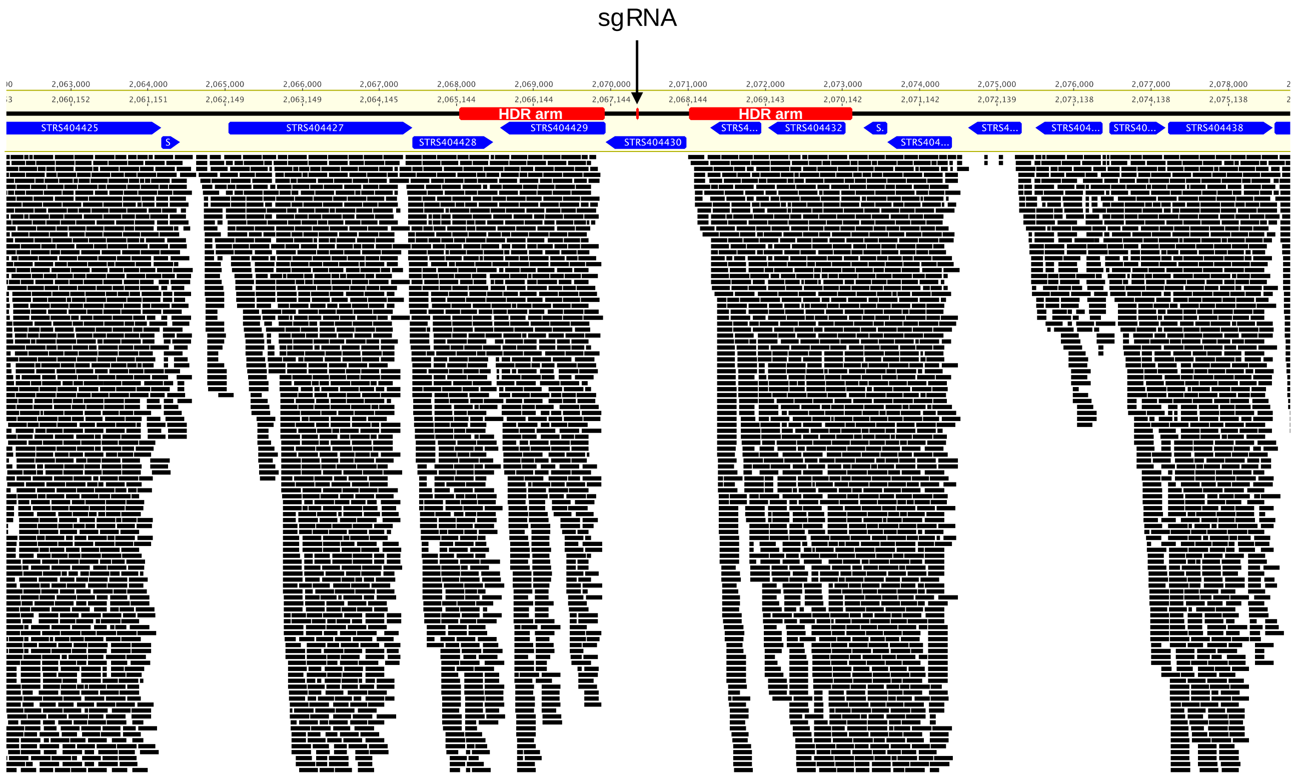


**Figure S4.** Mutagenesis of the albaflavenone BGC in *Streptomyces albus S4* Δ5. Black rectangles represent Illumina MiSeq reads mapped to the *S. albus* S4 Δ5 genome. Red rectangles indicate the homology-directed repair arms used to repair the double strand break generated by the Cas9 targeting the *STRS404430* gene. HDR, homology-directed repair arm; sgRNA, single guide RNA.

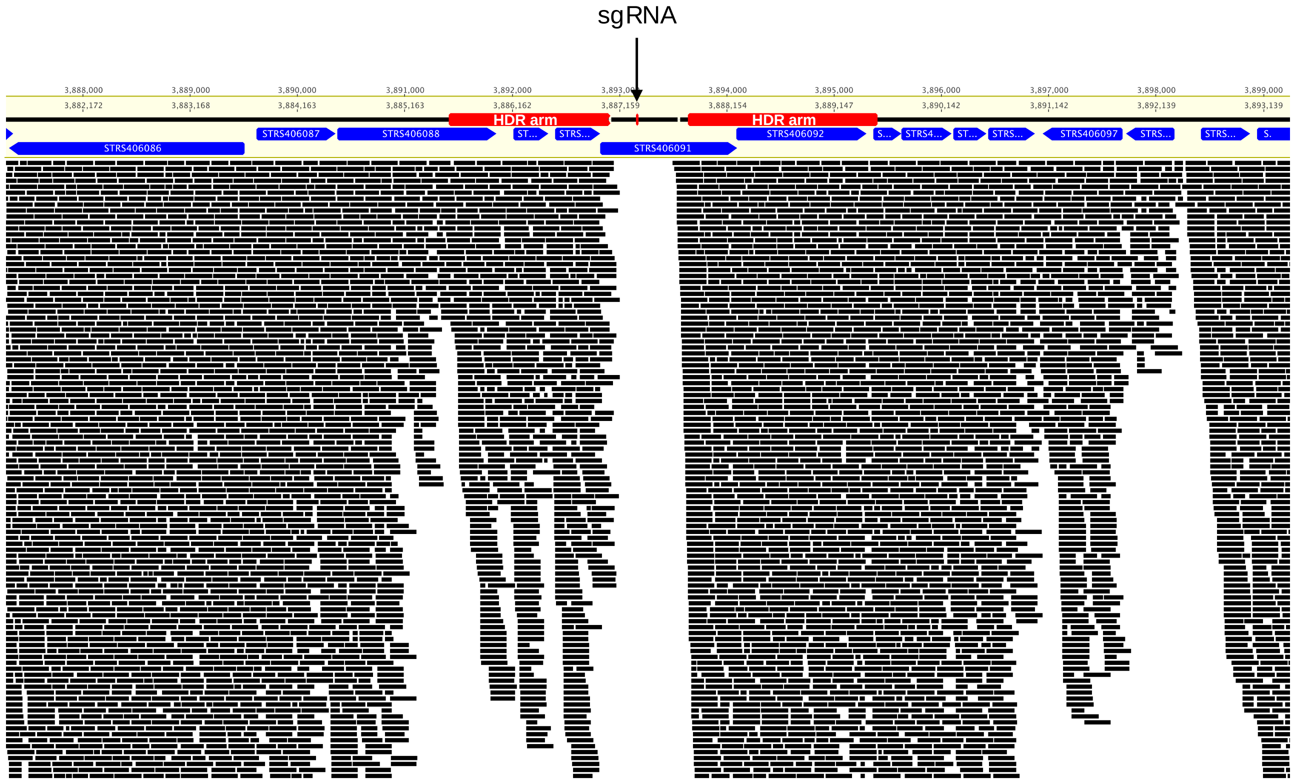


**Figure S5**. Mutagenesis of the fredericamycin BGC in *Streptomyces albus* S4 Δ5. Black rectangles represent Illumina MiSeq reads mapped to the *S. albus* S4 Δ5 genome. Red rectangles indicate the homology-directed repair arms used to repair the double strand break generated by the Cas9 targeting the *STRS406091* gene. HDR, homology-directed repair arm; sgRNA, single guide RNA.

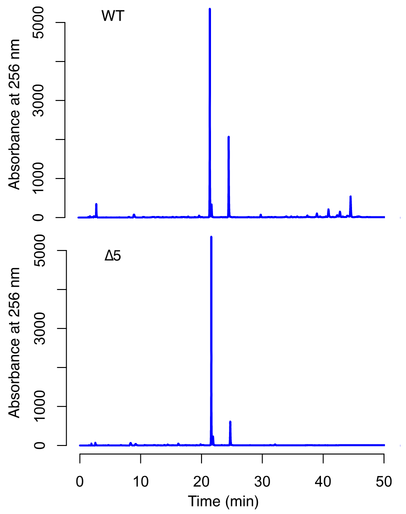


**Figure S6**. Full scale version of the HPLC chromatogram depicted in Figure 3 in the main text.

**Table S1.** Bacterial strains, cosmids and plasmids used in this study

| Strain/cosmid/plasmid | Description [a] | Reference |
|---|---|---|
| *Candida albicans* | *Candida albicans* strain CA-6 | [1] |
| *Micrococcus luteus* | *Micrococcus luteus* NCTC 9379 | NCTC |
| ***Streptomyces*** | | |
| S4 | Wild type *Streptomyces albus* S4 | [2] |
| Δ1 | S4 harbouring a complete deletion of the antimycin BGC | This study |
| Δ2 | Δ1 harbouring a deletion in the *STRS402234* (*fscC*) of the candicidin BGC | This study |
| Δ3 | Δ2 harbouring a deletion in the *STRS406449* gene of the surugamide BGC | This study |
| Δ4 | Δ3 harbouring a deletion in the *STRS404430* gene of the albaflavenone BGC | This study |
| Δ5 | Δ4 harbouring a deletion in the *STRS406091* gene of the fredericamycin BGC | This study |
| Δ5/Act | *S. albus* S4 Δ5 harbouring the actinorhodin BGC (*attB* ΦC31::pAH77); $Apr^R$ | This study |
| Δ5/Cin | *S. albus* S4 Δ5 harbouring the cinnamycin BGC (*attB* ΦC31:: pIJ10109) ; $Apr^R$ | This study |
| Δ5/Prun | *S. albus* S4 Δ5 harbouring the neoantimycin/prunustatin BGC split over two cosmids (*attB* ΦC31::cosmid813 *attB* ΦBT1::cosmid69); $Apr^R$, $Hyg^R$, $Kan^R$ | [3] |
| ***Escherichia coli*** | | |
| BW25113 | Host for REDIRECT PCR targeting system | [4] |
| NEBα | General cloning host | New England Biolabs |
| ET12567 | Non-methylating host for transfer of DNA into *Streptomyces* spp. (*dam, dcm, hsdM*); $Cam^R$ | [5] |
| GB05-red | Host for RecET recombination | [6] |
| **Cosmids and BACs** | | |
| Supercos1 | Cosmid backbone for *S. albus* S4 Cosmid 213; $Carb^R$, $Kan^R$ | Stratagene |
| Cosmid 213 | Supercos1 derivative containing the entire antimycin biosynthetic gene cluster; $Carb^R$, $Kan^R$ | [7] |
| Cosmid 213Δ*antF* | Cosmid 213 derivative harbouring an apramycin-marked deletion of *antF*; $Carb^R$, $Kan^{R,}$ $Apr^R$ | This study |
| Cosmid 213Δ*antA-antO* | Cosmid 213 derivative harbouring an apramycin-marked deletion in the entire antimycin biosynthetic gene cluster (*antABCDEFGHIJKLMNO*); *antF*; $Carb^R$, $Kan^{R,}$ $Apr^R$ | This study |
| Cosmid 213ΔantA-antO-FLP | Cosmid 213Δ*antA*-Δ*antO* derivative in which the apramycin resistance cassette was removed by the FLP recombinase; $Carb^R$, $Kan^R$ | This study |

| | | |
|---|---|---|
| Cosmid 213Δ*antA*-*antO*-FLPHygoriT | Cosmid 213Δ*antA*-*antO*-FLP derivative in which the *bla* resistance gene on the cosmid backbone was disrupted with a *hyg*$^{R}$-*oriT* cassette | This study |
| pIJ10109 | Derivative of pOJ436 harbouring the cinnamycin biosynthetic gene cluster from *Streptomyces cinnamoneus*; Carb$^{R}$, Apr$^{R}$ | [8] |
| Cosmid 813 | Supercos1 derivative harbouring *natABCDEF* from the neoantimycin biosynthetic gene cluster; Carb$^{R}$, Kan$^{R}$ | [3] |
| Cosmid 69 | Supercos1 derivative harbouring a partial *natB* gene and *natCDEFGQF'G'HIJKLNOP* genes from the neoantimycin biosynthetic gene cluster; Carb$^{R}$, Kan$^{R}$ | [3] |
| Cosmid 69- ΦBT1 | Cosmid 69 derivative engineered to integrate into the ΦBT1 *attB* site; Carb$^{R}$, Hyg$^{R}$ | [3] |
| **Plasmids** | | |
| pCRISPomyces-2 | pGM1190 derivative harbouring the CRISPR/Cas9 machinery; Apr$^{R}$ | [9] |
| pCRISPomyces-2-sur | Derivative of pCRISPomyces-2 derivative containing the *STRS406449*-targeting protospacer cloned into the BbsI site and homology-directed repair arms cloned into the XbaI site; Apr$^{R}$ | This study |
| pCRISPomyces-2-alb | Derivative of pCRISPomyces-2 derivative containing the *STRS404430*-targeting protospacer cloned into the BbsI site and homology-directed repair arms cloned into the XbaI site; Apr$^{R}$ | This study |
| pCRISPomyces-2-fdm | Derivative of pCRISPomyces-2 derivative containing the *STRS406091*-targeting protospacer cloned into the BbsI site and homology-directed repair arms cloned into the XbaI site; Apr$^{R}$ | This study |
| pIJ773 | ReDirect PCR template plasmid harbouring an apramycin resistance cassette and *oriT*; Carb$^{R}$, Apr$^{R}$ | [4] |
| pIJ10701 | ReDirect PCR template plasmid harbouring a hygromycin resistance cassette and *oriT*; Hyg$^{R}$ | [4] |
| pKC1132-UpDn | Derivative of suicide plasmid pKC1132 [10]<br>containing ~3kb of homologous DNA upstream and downstream of the region of *fscC* targeted for deletion; Apr$^{R}$ | [11] |
| pUZ8002 | Encodes conjugation machinery for mobilisation of plasmids from *E. coli* to *Streptomyces;* Kan$^{R}$ | [5] |

[a] Carb, carbenicillin; Apr, apramycin; Hyg, hygromycin, Kan, kanamycin; Cam, chloramphenicol; *oriT*, origin of conjugal transfer

**Table S2.** Oligonucleotide primers used in this study

| Primer alias | Sequence (5'-3')[a] | Description |
|---|---|---|
| RFS196 | **ctcgtgtcgttctcaggtggagaggtgcctgcgcgctca**tgtaggc tggagctgcttc | PCR: *antF* REDIRECT knockout cassette |
| RFS236 | cgcctacaacaccggtgagt | PCR: confirmation of Δant mutation |
| RFS237 | aggggacgatgttgacgacc | PCR: confirmation of Δant mutation |
| RFS197 | **gacggccccggcggccgggacggccggcggtgcctgatg**attcc ggggatccgtcgacc | PCR: *antF* REDIRECT knockout cassette |
| RFS219 | **cacgcgcccgcgtgtctcaccccgccatggtggccgtca**attccg gggatccgtcgacc | PCR: *antABCDEHIJKLMNO* REDIRECT knockout cassette |
| RFS203 | **ccgcctcggccgggtcgggagacatctggcgggcggtca**tgtag gctggagctgcttc | PCR: *antABCDEHIJKLMNO* REDIRECT knockout cassette |
| RFS242 | atcacgcggctgatcgacca | PCR: confirmation of Δ*antF* mutant strain |
| RFS243 | tggaggaactgcggaccatc | PCR: confirmation of Δ*antF* mutant strain |
| EH_S3 | acgctgccgggccgccgcgagaaa | CRISPR protospacer targeting *STRS4_04430* (albaflavenone BGC) |
| EH_S4 | aaactttctcgcggcggcccggca | CRISPR protospacer targeting *STRS4_04430* (albaflavenone BGC) |
| EH_S7 | acgcgtacgcctgctccatggaga | CRISPR protospacer targeting *STRS4_06091* (fredericamycin BGC) |
| EH_S8 | aaactctccatggagcaggcgtac | CRISPR protospacer targeting *STRS4_06091* (fredericamycin BGC) |
| EH_S9 | acgccacctcacgcggcaccggga | CRISPR protospacer targeting *STRS4_06449* (surugamide BGC) |
| EH_S10 | aaactcccggtgccgcgtgaggtg | CRISPR protospacer targeting *STRS4_06449* (surugamide BGC) |
| EH_P7 | **tgccgccgggcgttttttat**gtgtactggttccgctc | PCR: *STRS4_04430* homology-directed repair arm |
| EH_P8 | **tttgttcgtgcctgctttcc**cgaatccaccaccgaac | PCR: *STRS4_04430* homology-directed repair arm |
| EH_P9 | **ggcgttcggtggtggattcg**ggaaagcaggcacgaac | PCR: *STRS4_04430* homology-directed repair arm |
| EH_P10 | **cggcctttttacggttcctggcct**ggttggatgcgaagagg | PCR: *STRS4_04430* homology-directed repair arm |
| EH_P15 | **tgccgccgggcgttttttat**cgctcttcctgctgttgttgg | PCR: *STRS4_06091* homology-directed repair arm |
| EH_P16 | **ggtcatgtggtagccgttgg**ttggcccttcacgatcatctcc | PCR: *STRS4_06091* homology-directed repair arm |
| EH_P17 | **agatgatcgtgaagggccaa**ccaacggctaccacatgacc | PCR: *STRS4_06091* homology-directed repair arm |
| EH_P18 | **cggcctttttacggttcctggcc**tatggtcttgaggtcgttgaagc | PCR: *STRS4_06091* homology-directed repair arm |
| EH_P19 | **tgccgccgggcgttttttat**gtacgtcatgtccacctcc | PCR: *STRS4_06449* homology-directed repair arm |
| EH_P31 | **cgtaagcctgggtgttgtcc**tacagctcgctcagttcg | PCR: *STRS4_06449* homology-directed repair arm |
| EH_P27 | **cgcgaactgagcgagctgta**ggacaacacccaggcttacg | PCR: *STRS4_06449* homology-directed repair arm |
| EH_P28 | **cggcctttttacggttcctggcct**ctccttcaccgacttcagc | PCR: *STRS4_06449* homology-directed repair arm |

| | | |
|---|---|---|
| EH_P35 | gtcgtgaatctcctgatcg | PCR: confirmation of *STRS4_04430* mutation |
| EH_P36 | tacggctacctctacatcgacc | PCR: confirmation of *STRS4_04430* mutation |
| EH_P37 | tggccgaacccttctactcc | PCR: confirmation of *STRS4_06091* mutation |
| EH_P38 | ccatgtccaggtcgttcagc | PCR: confirmation of *STRS4_06091* mutation |
| EH_P39 | caccaggacttcttcacg | PCR: confirmation of *STRS4_06449* mutation |
| EH_P40 | gagggagaagaagttgtcgtgg | PCR: confirmation of *STRS4_06449* mutation |

[a] non-homologous sequences are underlined and engineered restriction endonuclease sites are bolded
